# Spatially resolving how phosphorylation affects ß-cardiac myosin activity in porcine myofibril sarcomeres with single molecule resolution

**DOI:** 10.1101/2024.06.27.600847

**Authors:** Matvey Pilagov, Sonette Steczina, Ateeqa Naim, Michael Regnier, Michael A. Geeves, Neil M. Kad

## Abstract

Cardiac muscle contraction is mediated by myosin binding from the thick filament of the sarcomere to the thin filament in an ATP powered reaction. This process is highly regulated on a beat-to-beat basis by calcium interactions with the thin filament. Additionally, the number of heads available for participation in contraction is also regulated, resulting in a dynamically variable reserve of heads for controlling contractile force. We aimed to discover the size of this reserve and how it is modulated by phosphorylation. Using single molecule imaging of fluorescently labelled ATP molecules binding and releasing myosins within porcine cardiac sarcomeres, we could determine myosin activity with high spatial resolution. We find three kinetic species when examining the myosin ATPase. The fastest is consistent with non-specific ATP binding to myosin’s surface, and the slower two species are consistent with the previously identified DRX and SRX states. The former is thought to represent myosins in an ON state, ready to interact with the thin filament and the latter an OFF state with slowed ATPase that constitutes the cardiac reserve. We find that the cardiac reserve is 50% in the sarcomere and this can be sub-divided into the P-, C- and D-zones, with the D-zone having the least population of OFF heads (44%). Treatment with PKA phosphorylates cardiac myosin binding protein-C (cMyBP-C) leading to a 16% reduction in reserve in the C-zone (where cMyBP-C is found), a 10% reduction in the P-zone, and an unexpected 8% increase in the D-zone. By contrast, myosin regulatory light chain (RLC) phosphorylation with myosin light chain kinase (MLCK) resulted in a large 24% decrease in reserve myosins, interestingly the least affected area of the sarcomere was the C-zone. Altogether these data suggest that cMyBP-C’s interaction with RLC governs the degree of activation due to RLC phosphorylation.

## Introduction

Cardiac muscle contraction is dynamically regulated to ensure rapid and constant adaptation to the changing physiological demands placed on the heart. The fundamental contractile organelle in cardiac muscle is the myofibril, which is comprised of a tandem array of sarcomeres. Sarcomeres consist of myosin containing thick filaments and actin containing thin filaments, both of which are regulated to control the timing and force of contraction. Regulation of thin filaments is mediated by calcium binding to thin filament associated control proteins and has been well characterized (Gordon, Homsher, and Regnier 2000), although some questions remain. However, our understanding of thick filament regulation is still being developed. The current view of thick filament regulation is that myosins adopt two biochemical states under relaxed conditions, one poised to interact with actin termed the disordered relaxed (DRX) state, and a second super-relaxed (SRX) state that corrals a reserve population of myosins in a very depressed activity state (Stewart et al. 2010; Hooijman, Stewart, and Cooke 2011; Nogara et al. 2016). This reserve enables the heart to recruit myosins for stronger contraction, but to save energy when they are not immediately required.

Recent high-resolution structural studies have provided valuable insights into the complex environment of the sarcomere (Tamborrini et al. 2023; Nag et al. 2023). These complement classical studies (R. Craig and Offer 1976; Tonino et al. 2019; Bennett and Gautel 1996) that have used counter-staining to define regions of the thick filament based on their protein composition. The P-zone is closest to the sarcomere center (M-line) and possesses myosin binding protein-H (MyBP-H), the C-zone is marked by the clear presence of 9 bands of myosin binding protein-C (MyBP-C) and the D-zone accounts for the remainder - occupying approximately the last third of the thick filament. Interestingly, despite the use of compounds thought to promote myosin into a stabilized, folded-back (interacting heads motif (IHM)) state, the D-zone was not resolved in these structural studies. This suggests that the ancillary myosin binding proteins could assist in generating order on the thick filament backbone. Myosin’s biochemical activity is mediated by load (Linari et al. 2015; Fusi et al. 2016), by direct calcium binding to both thin and thick filaments (Ma et al. 2023; Lehman and Kendrick-Jones 1973; Bremel and Weber 1975; Z. A. Podlubnaya, Malyshev, et al. 2000; Z. Podlubnaya et al. 1999; Z. A. Podlubnaya, Kąkol, et al. 2000) , and by phosphorylation (Heling, Geeves, and Kad 2020; Colson et al. 2010; Levine et al. 1996; Ponnam et al. 2019; McNamara, Singh, and Sadayappan 2019). Phosphorylation affects a number of proteins in the sarcomere, however, two in particular alter myosin’s activity. Myosin’s regulatory light chain (RLC) is constitutively phosphorylated at a ∼40% level, suggesting only one of each of myosin’s two heads is phosphorylated at any time (Colson et al. 2010; Scruggs et al. 2010; Chang et al. 2016). However, further phosphorylation activates myosins by pushing the heads away from the thick filament backbone (Colson et al. 2010; Levine et al. 1996). This phosphorylation is mediated by myosin light chain kinase (MLCK), a calmodulin dependent PKC-induced enzyme. By contrast, the ß-adrenergic response pathway activates protein kinase-A (PKA), which rapidly phosphorylates sites on MyBP-C and results in the de-repression of myosin activity (Colson et al. 2010; Ponnam et al. 2019; Tong et al. 2008; Previs et al. 2016). Through these mechanisms of phosphorylation, the ratio of SRX and DRX myosins in the myofibril can be altered, which is crucial to changing the force that can be generated. Both the dynamics of the SRX-DRX equilibrium (Mohran et al. 2024) and the distribution of the SRX/DRX states along the thick filament remain unclear. Consistent with MyBP-C affecting this ratio (McNamara, Singh, and Sadayappan 2019; McNamara et al. 2016), recent attempts to localize the distribution of SRX and DRX myosins within the sarcomere have shown that the SRX state predominates in the C-zone of skeletal muscle (S. R. Nelson et al. 2020; Pilagov et al. 2023) and in α-cardiac myosin in mice (S. Nelson et al. 2023). The pathophysiological importance of understanding how phosphorylation affects activity is highlighted by the observation that cardiac myosin binding protein-C (cMyBP-C) is one of the two most prevalent locations for mutations associated with hypertrophic cardiomyopathy (Konno et al. 2010), and RLC mutations are also (although much less frequently) associated with the same disease (Poetter et al. 1996; Maron and Maron 2013).

Here we sought to understand the effects of phosphorylation on myosin recruitment in a human analogous ß-cardiac porcine background. To do this, we applied a single molecule approach to detect the ATPase activity of predominantly ß-cardiac myosin from porcine left ventricular cardiac myofibrils with sub-sarcomeric resolution. Untreated myofibrils directly extracted from tissue showed SRX myosin population of ∼50%, distributed as 57%, 54% and 44% across the P-, C- and D-zones respectively. Upon treatment with PKA, this changed to 47%, 37% and 52% (P-, C- and D-zones respectively) indicating a strong zonal difference explained by cMyBP-C phosphorylation in the C-zone. Using MLCK to phosphorylate RLC, we observed a global increase in the DRX myosin population, with 30%, 40% and 19% SRX in the P-, C- and D-zones respectively. The reduced level of activation in the C-zone may suggest an interaction between cMyBP-C and RLC, which limits the amount of myosin that can be made available for interaction with actin.

## Methods

No live animals were used in these studies. Muscle tissue was collected in accordance with the U.K. Animals (Scientific Procedures) Act 1986 and associated guidelines.

### Dissection and storage of porcine heart

Immediately following excision, the heart of a freshly euthanized adult farm pig was submerged in ice-cold cardioplegic solution (5.5 mM Glucose, 0.5 mM MgSO_4_, 24 mM KCl, 20 mM NaHCO_3_, 109 mM NaCl, 0.9 mM H_2_NaO_4_P, 1.8 mM CaCl_2_, 0.01% NaN_3_, pH 7.4). While remaining in solution, left ventricular trabeculae samples were extracted from the heart and cut into ∼5 mm thick pieces, immediately flash frozen, and kept at -80°C for long term storage.

### Myofibril Isolation and fluorescent staining

Myofibril isolations were carried out as previously described in Pilagov et al., 2023 with adaptations for cardiac tissue. Briefly, a single porcine left ventricular trabeculae sample was rapidly thawed in chilled Prep Buffer (20 mM MOPS, 132 mM NaCl, 5 mM KCl, 4 mM MgCl_2_, 5 mM EGTA, 10 mM NaN_3_, 5 mM DTT, 20 mM 2,3-Butanedione monoxime (BDM), protease inhibitor cocktail (A32965; Thermo Scientific), pH 7.1) (Vikhorev, Ferenczi, and Marston 2016) and cut into 2-3 mm thick strips. Samples were secured to a Sylgard^TM^ PDMS-based petri dish with tungsten rods at both ends of the tissue and Prep buffer was replaced with Permeabilization buffer (Prep buffer + 0.5% Triton X-100). Samples were permeabilized overnight with gentle agitation (4°C) and then rinsed three times with Prep buffer to remove Triton X-100. Permeabilized samples were cut down to ∼1 mm thick strips, then homogenized in Prep buffer using a Tissue Ruptor II at the lowest speed for 10 seconds twice with a 1-minute rest period on ice in between. The concentration of myofibril preparations was kept constant by targeting an optical density (OD) of ∼0.4 at 600 nm at the end of the preparation.

For fluorescent staining of Z-discs, isolated myofibrils were incubated for 1.5 hrs at 4°C on a rotator with 11 nM anti-α-actinin mouse antibody (A7811; Sigma-Aldrich), 5.5 nM Alexa-488 goat anti-mouse IgG (A11001; ThermoFisher Scientific) and 1 mg/ml bovine serum albumin (BSA), to avoid non-specific antibody binding.

### Imaging Conditions

Microfluid imaging chambers were constructed and coated with 15 µg/ml ≥300KDa Poly-L-Lysine (PLL, Sigma) on the day of imaging as described previously (Pilagov et al. 2023) and rinsed with 100 µl Prep buffer. Fluorescently tagged myofibrils were pipetted into the imaging chamber and incubated for 30 min at 4°C to allow adhesion of myofibrils to the PLL coated surface. Excess non-adherent myofibrils were flushed out with two washes of 100 µl No BDM Buffer (Prep Buffer without BDM) with a 1 min rest at 4°C in between. Finally, 100 µl Imaging Buffer (No BDM Buffer plus 1-10 nM Cy3-ATP, 3.27 mM ATP, and 2 mM Trolox) was added to the flow chamber and allowed to equilibrate for 10 min prior to imaging. Cy3-ATP was synthesized and kindly provided by Dr. C.P. Toseland (University of Sheffield, Sheffield, UK) (Toseland and Webb 2011). To ensure no Cy3-ADP contamination, the nucleotide was enzymatically regenerated into Cy3-ATP prior to the experiments, as described previously (Pilagov et al. 2023). All solutions were gently flowed into the imaging chamber at ∼100 µl/min to prevent dislodging of myofibrils from the surface.

### Kinase Treatment

For this study, a subset of myofibrils were pre-treated with PKA or MLCK. Incubations were carried out on a rotator for constant agitation. PKA (cAMP-dependent Protein Kinase (PKA), catalytic subunit, New England Biolabs, P600S) treatment at 62.5 units/µl was used with the recommended buffer (NEBuffer™ for Protein Kinases (PK) + 200 µM ATP) at 4°C for 1 hour. Recombinant human smooth muscle Myosin light chain kinase (MLCK, Abcam, ab55674) has previously been shown to phosphorylate cardiac RLC (cRLC (Sevrieva et al. 2020; Turner et al. 2023; Park-Holohan et al. 2021)) and was used at 0.4 µM, following the previously described protocol (5 mM BDM, 67 µM CaCl_2_, 6 µM Calmodulin) (Stelzer, Patel, and Moss 2006). The reaction was carried out at 4°C for 1 hour. All enzymes were incubated at lower temperatures for longer periods of time than recommended to prevent myofibril degradation.

### Image acquisition

Images were acquired at 21°C on a custom-built oblique angle fluorescent (OAF) microscope as described previously (Desai, Geeves, and Kad 2015). For excitation of the Cy3-ATP, a 561 nm diode OBIS LS laser (Coherent, USA) was used at 20 mW and the Alexa 488-anti-α-actinin was excited with an Oxxius 488-nm laser at 18 mW. To prevent fluorescence photobleaching, myofibrils were located using brightfield. Photobleaching was also ameliorated during image acquisition via the use of stroboscopic illumination as previously described (Pilagov et al. 2023). Stroboscopic illumination resulted in a six-frame repeat pattern consisting of five sequential, 200 ms 561 nm laser Cy3-ATP frames with a dark period of 1800 ms. The sixth frame was a 200 ms 488 nm illumination during the 1800 ms dark period of the fifth 561 nm laser cycle to capture an image of the Z-disc. Each sample was imaged for 30 minutes producing 1000 frame videos comprised of the Z-disc and Cy3-ATP frames. To prevent ADP and Cy3-ADP accumulation, chambers were used for no longer than one hour.

### Data Analysis

To extract the spatiotemporal data of Cy3-ATP binding events, analysis was carried out using a series of custom-written scripts. Firstly, a Python script divided the 1000 frame video into two videos, one containing all the ATP frames and the other containing all the Z-disc frames. TrackMate, an ImageJ plugin (Tinevez et al. 2017; Ershov et al. 2022), was utilized for automated tracking and super-localization of fluorescent Cy3-ATP at the Z-discs. The following parameters were used to run TrackMate: estimated object diameter for the Laplacian of Gaussian (LoG) detector: 390 nm, linking and gap-closing distances: 100 nm, and gap-closing max frame: 5 frames. All videos were manually checked against the TrackMate output to ensure correct assignment of the tracks. If myofibril drift was identified across the duration of a video acquisition, this was corrected using a star-map approach (https://github.com/fiji/Descriptor_based_registration) prior to event tracking. These scripts are available and annotated (see data availability statement).

TrackMate provided super-localized x- and y-positions of Cy3-ATP events and their durations, which were then extracted to Microsoft Excel for further analysis. Based on our previously described method (Pilagov et al. 2023) we corrected any tilt using Z-disc images and then used these as fiducial markers to resolve Cy3-ATP positions to zones of the A-band. We calculated the precision for both Z-disc (X - 68.2 nm, Y – 135.4 nm) and Cy3-ATP (X - 28.1 nm, Y – 38 nm) events (Gelles, Schnapp, and Sheetz 1988). Cy3-ATP events were plotted as cumulative residence time histograms for both zonally separated events (P-, C-, and D-zones) and events across the whole thick filament (denoted as “all zones” data). Cumulative residence time histograms were fitted to three exponentials, with smaller and larger numbers of exponentials tested to ensure the optimal number was chosen for the final analysis. For accurate fitting within the sub-sarcomeric zones, we used two constraints based on our fitting of the ‘all zones’ data: the sum of the number of zonal events were constrained within 20% of that which was determined for the ‘all zones’ data and the rate constants per zone were permitted to vary within 20% of the ‘all zone’ rate constants. Based on our previous study (Pilagov et al. 2023), the three fitted exponentials corresponded to: 1) non-specific association of Cy3-ATP with myosin (Ušaj et al. 2021; Amrute-Nayak et al. 2014), 2) Cy3-ATP turnover association with myosin in the DRX state and, 3) Cy3-ATP turnover association with myosin in the SRX biochemical state. A visual breakdown of three components of a typical fit can be seen in Fig.1D.

**Figure 1.**
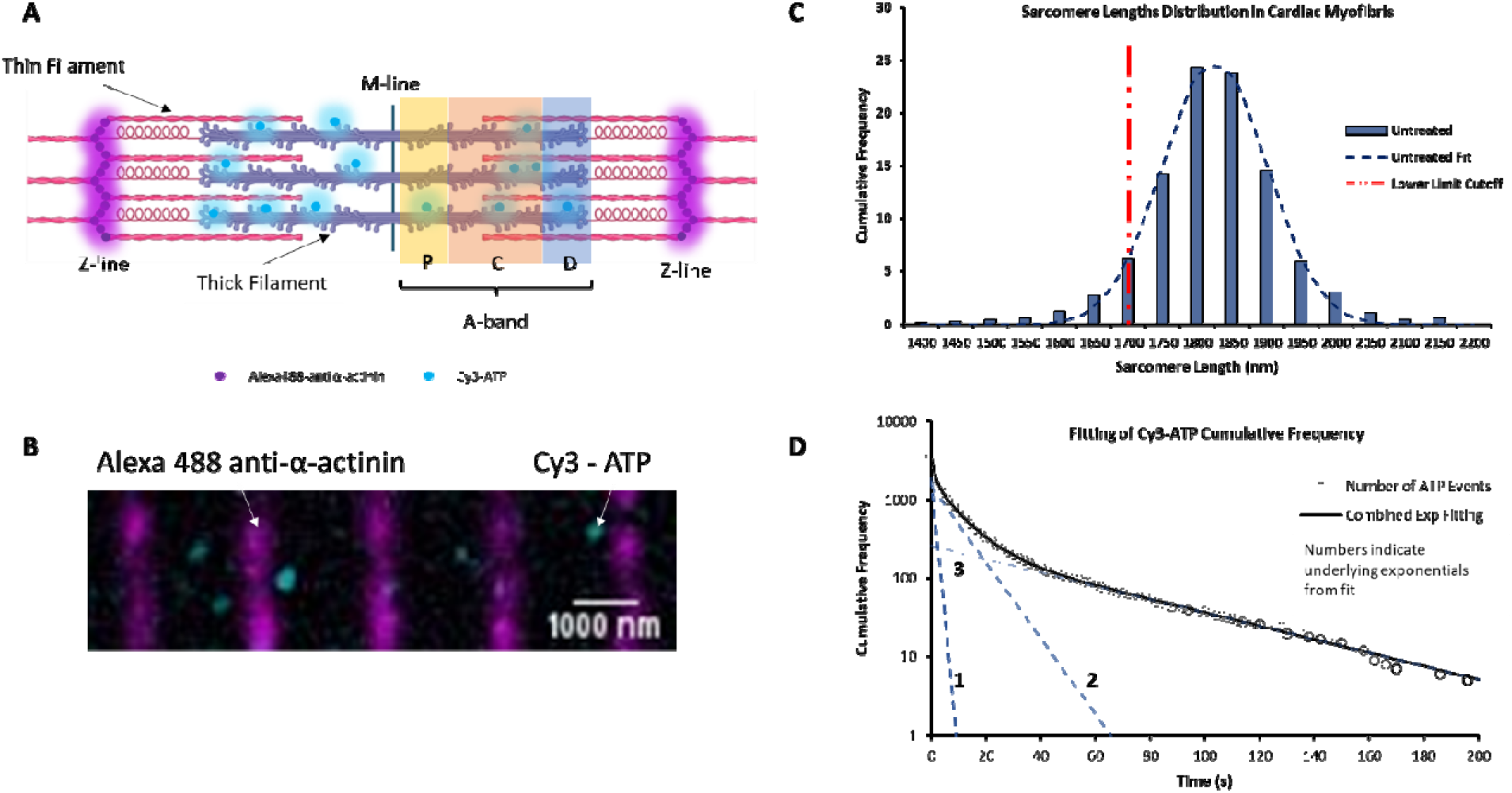
Single molecule tracking of fluorescently labeled ATP turnover in porcine cardiac myofibrils. **A)** A schematic representation of a single sarcomere. P-, C-, and D-zones of the thick filament are labelled accordingly. Alexa 488 – labelled α-actinin is denoted with magenta circles, while Cy3-ATP binding to the thick filament is represented as cyan circles. **B)** An enlarged section of a representative porcine cardiac left ventricular myofibril as observed following the overlay of 561 nm illuminated Cy3-ATP (cyan) and 488 nm illuminated α-actinin containing z-discs (magenta). **C)** Distribution of sarcomere length for untreated myofibrils (N=3 pigs, n=647 sarcomeres) at 21°C. Distribution of sarcomere lengths for other conditions can be found in Figure S1. The red dotted line indicates the threshold for the smallest sarcomere that we analyzed for all our datasets. **D)** Cumulative residence time histogram showing the distribution of ATP attachment event durations in untreated porcine cardiac myofibrils (unfilled black circles; N=3 pigs, n=3638 events). Dashed lines show individual exponential fits (numbered 1 to 3 fastest to slowest) that make up a cumulative fit of all three exponentials (black solid line). Note the frequency axis is plotted on a logarithmic scale.

### Phosphoprotein Gels

Tris-Glycine gels (gel percentages given in figure legends) were either self-casted or purchased from ThermoFisher (XP04205BOX) and the ladder used was PageRuler™ Plus Prestained Protein Ladder (ThermoFisher 26619). Samples were boiled at 95 °C for 5 minutes with standard SDS Sample Loading Buffer prior to loading. Gels were stained with Pro-Q Diamond Phosphoprotein Gel Stain (ThermoFisher Scientific, P33300) to determine the phosphorylation levels of sarcomeric proteins. Gels were subsequently stained with Coomassie gel dye to quantify total protein per sample. All data were calculated as relative changes in phosphorylation normalized to loading using the same band from the Coomassie gel. Pro Q gels were imaged using Syngene G:BOX Chemi XX6 set to Pro-Q imaging setting (green lamp, 605 nm filter) and Coomassie was imaged using the white light setting.

### Statistics

Accuracy for TrackMate super-resolved events for both Cy3-ATP events and Z-discs were established using variance from fixed points in the image as described previously (Gelles, Schnapp, and Sheetz 1988; Pilagov et al. 2023). Errors for cumulative residence time histogram fits were determined using SolverStat (Comuzzi et al. 2003) and were expressed as standard error of the mean (SEM) with n as the number of Cy3-ATP attachments for the respective zone. Errors were propagated through calculations using standard propagation formulae (Caldwell and Vahidsafa 2022). Significance analysis was performed using a Student T-test within each zone. A threshold for statistical significance was set as p<0.05 and is indicated by asterisks (*). For this paper, N equals the number of porcine farm pigs used as biological replicates.

## Results

### Imaging fluorescent ATP lifetimes in relaxed myofibrils

The sarcomere consists of interdigitated thick and thin filaments flanked by α-actinin containing Z-discs (Fig 1A). To determine where in the relaxed sarcomere myosin is actively turning over ATP, we first imaged the positions of the Z-discs as fiducial markers (Fig 1A, B). This was achieved using a fluorescently tagged anti α-actinin containing antibody (Fig 1B, magenta). From such images, we measured sarcomere length by using super-localization analysis (see below); this information was used to select for myofibrils with a sarcomere length above 1.7 µm (Fig 1C). This ensured the we used myofibrils in a clearly relaxed state for further analysis.

To determine the activity of individual myosins, we measured attachment durations of Cy3 fluorescently labelled ATP (Cy3-ATP), previously shown to have no impact on myosin’s activity (Pilagov et al. 2023; Toseland and Webb 2011; Ušaj et al. 2021). A ∼1e^6^ fold excess of non-fluorescent ATP (3.27 mM) was used to ensure that the myofibril remained relaxed throughout the experimental duration. However, given that observation times were ∼30 mins, it was necessary to include on-going corrections to the positions of the fiducial markers. Therefore, for every five Cy3-ATP imaging frames, an image of the Z-discs was taken to realign the video. During each Cy3-ATP frame (Fig 1B), the position of every fluorescent ATP was tracked and super-localized using the ImageJ plugin TrackMate. The attachment duration for each bound Cy3-ATP was derived from the video analysis to enable the assignment of Cy3-ATP’s bound-lifetime to its location within the sarcomeric A-band. Using the whole thick filament as a starting point, we binned all the lifetimes together as a cumulative residence time histogram (Fig 1D, ‘all zones’ data). As with our previous observations in skeletal muscle, (Pilagov et al. 2023) three populations best described the observed decay. These three populations fitted to lifetimes consistent with non-specific ATP binding (1.2 s; described in (Ušaj et al. 2021; Amrute-Nayak et al. 2014)), the DRX state (10.2 s) and the SRX state (155.9 s); these lifetimes were photobleach corrected as described previously (Pilagov et al. 2023).

### Phosphorylation affects the distribution of SRX and DRX in porcine myofibrils

Using the lifetimes to identify the populations of myosin activity, we were able to calculate the percent population of SRX myosins in the sarcomere. The amplitudes of each phase provided the starting point for estimating population. During single molecule imaging, faster events will be observed more frequently than slow turnover events, therefore, amplitudes were scaled against the observed rate constant to correct for this rate-induced artifact. To achieve this, the ratio of the rate constants (DRX/SRX) was multiplied by the SRX amplitude to calculate a corrected SRX population as previously described (Pilagov et al. 2023). Once corrected, the percent population of SRX for untreated myofibrils was 49.8% ± 0.03% across the thick filament (the non-specific population of ATP events was not included in these calculations). To identify if there were differences in the distribution of DRX and SRX myosins along the thick filament, we binned the super-resolution information provided by TrackMate into the P-, C- and D-zones. We observed similar percent populations of SRX in the P- and C-zones (56.8% ± 1.1% vs 53.8% ± 0.4%, respectively), however, the D-zone had a lower percentage of myosins in the SRX state (43.9% ± 0.5%). These results highlight that a more nuanced view of myosin activity within the sarcomere is critical for studies of the thick filament.

With an understanding of the untreated porcine cardiac myofibril established, we then used PKA to reproduce the effects of β-adrenergic stimulation of cardiac muscle. The main targets for PKA are cMyBP-C, cTnI, and Titin (Ahmed and Lindsey 2009; Krüger and Linke 2006). Using the Pro-Q phosphorylation gel stain, treatment with PKA led to a 2.5-fold increase in phosphorylation of cMyBP-C (Fig 2 B & C). cTnI was also phosphorylated, however, because cTnI does not impact the system in relaxed conditions we did not study it further. Examining the percent SRX using Cy3-ATP imaging revealed PKA phosphorylation had little effect on myosin activity when measured across all zones (46.1% ± 0.6% vs 49.8% ± 0.03% for untreated; Fig 3 and Table 1). However, more detailed analysis of the distribution of myosin activity within the sarcomere thick filament indicated, relative to untreated myofibrils, the SRX population after PKA phosphorylation was reduced by 16.4% in the C-zone (to 37.4% ± 4.7%, Fig 3 and Table 1), consistent with PKA phosphorylating cMyBP-C and de-repressing myosin activity. A smaller, 9.5% reduction in SRX population was also seen in the P-zone (to 47.3% ± 8.6%, Fig 3 and Table 1), possibly suggesting a cooperative effect from the C-zone. Surprisingly, the population of SRX increased by 8.3% in the D-zone (to 52.2% ± 1.4%, Fig 3 and Table 1), compensating for the decreases in the P- and C-zones. These results aid in explaining the lack of change in the SRX population across the whole sarcomere between untreated and PKA phosphorylated myofibrils.

**Figure 2.**
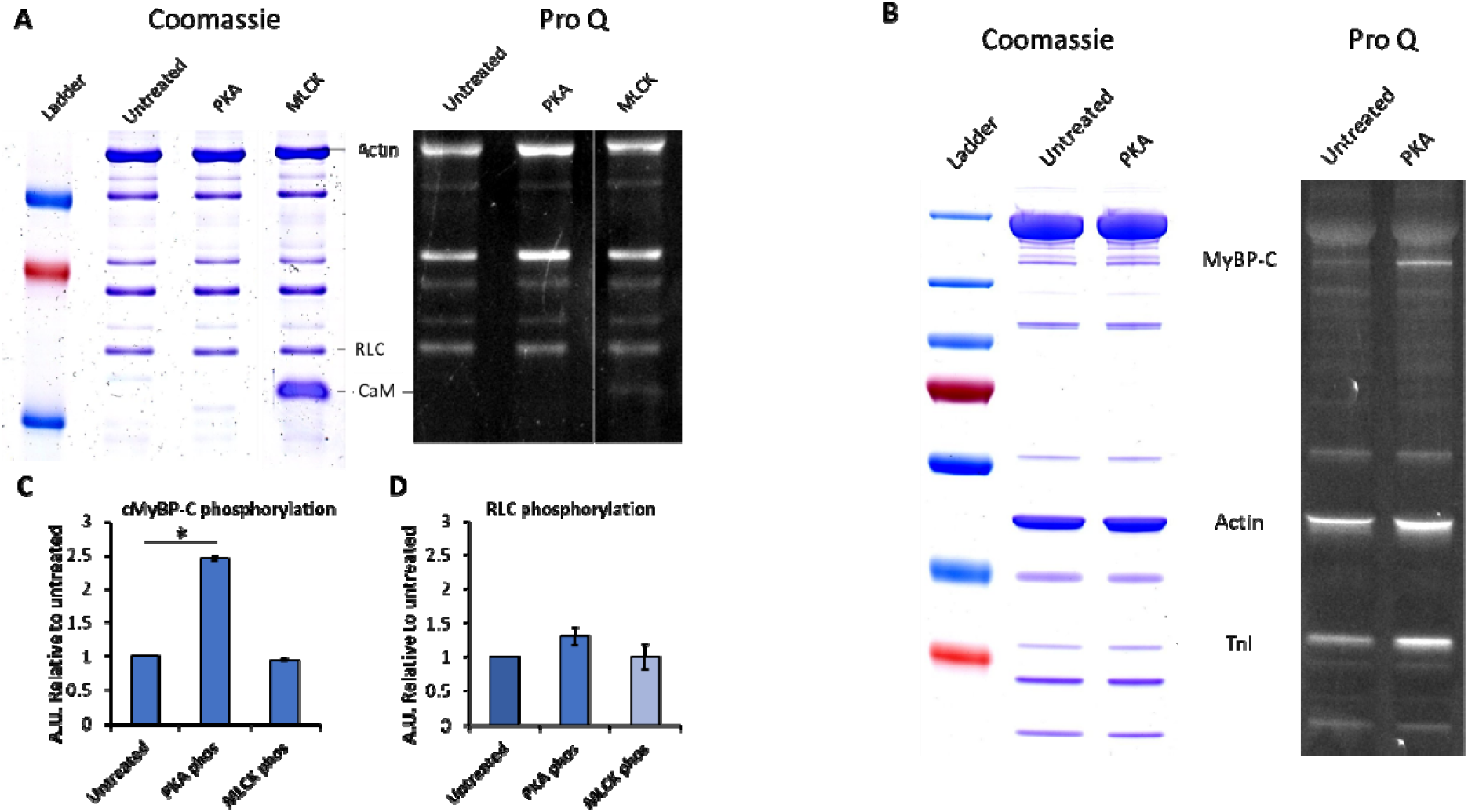
Quantification of cMyBP-C and RLC phosphorylation after PKA and MLCK treatment of porcine cardiac myofibrils. Protein was isolated from homogenized myofibrils and treated with PKA and MLCK and compared to untreated myofibrils. Myofibrils per condition were run on a 15% SDS PAGE gel (A) as well as a 4-20% gradient SDS-PAGE gel (B) to allow for better separation and resolution of the cMyBP-C band. Each gel was stained with Pro-Q Dimond Phosphoprotein Gel Stain (right) followed by Coomassie Blue stain (left) to quantify total protein. Densitometry analysis was performed on the cMyBP-C and RLC phosphorylation bands and normalized to the actin band of the respective Coomassie gel stain. (C) cMyBP-C phosphorylation per treatment group was calculated relative to the untreated group (N=1, n=2). (D) RLC phosphorylation per treatment group was calculated relative to the untreated group (N=1, n=3). Data shown as mean ± SEM. Student’s T-test was performed relative to untreated *p<0.05. Uncropped gels are provided in the supplementary material (Figure S2).

**Figure 3.**
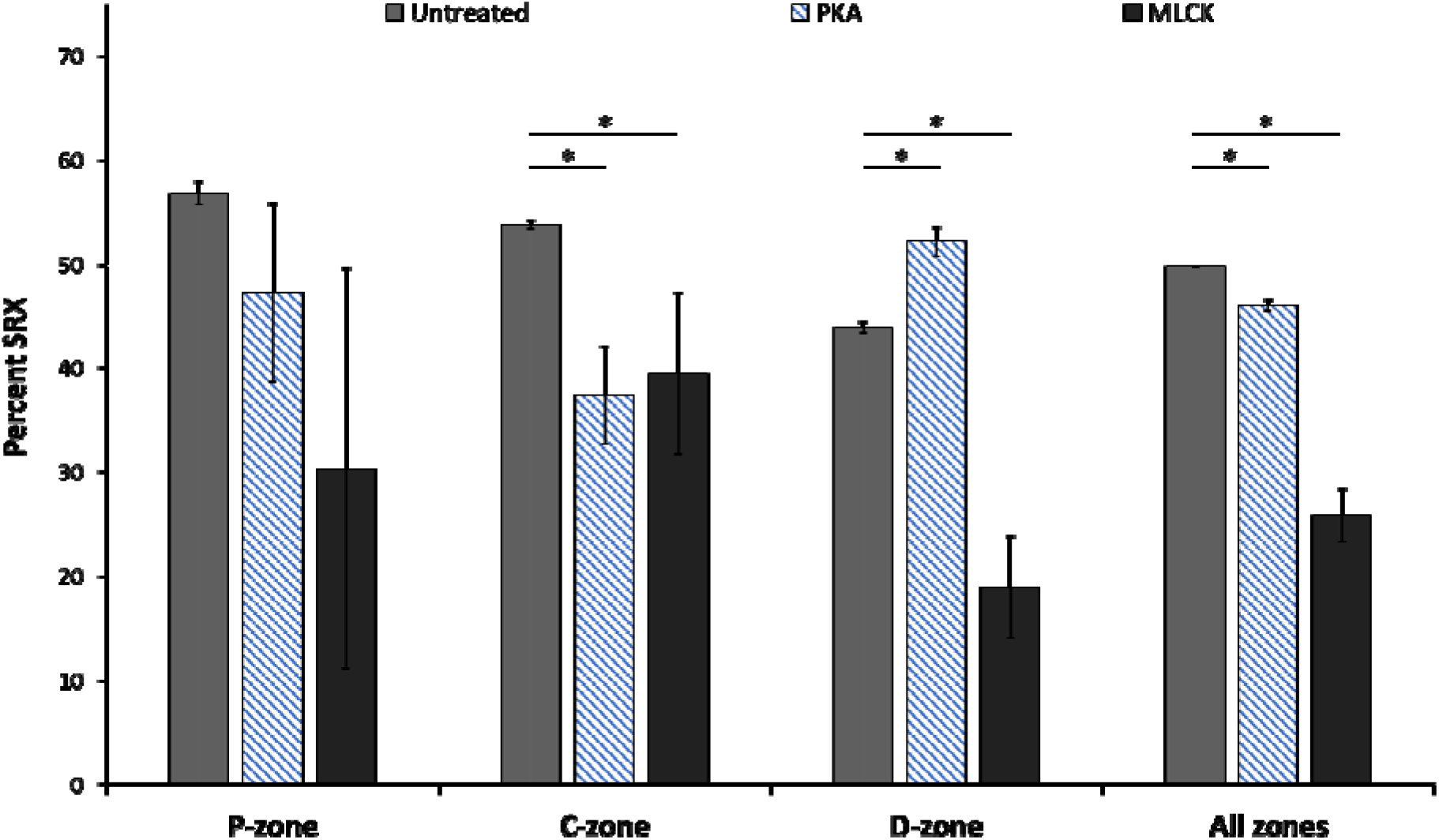
Differential impacts of cMyBP-C and RLC phosphorylation on the population of SRX myosins within the thick filament zones of porcine cardiac myofibrils. Cy3-ATP turnover rates were calculated as an average across all zones of the sarcomere as well as within the P-, C-, and D-zones. The Cy3-ATP turnover rates per zone were used to calculate the relative population amplitude of SRX myosin heads based on three exponential fits. Data was collected from untreated myofibrils (for N’s see Fig.1) or myofibrils treated with 62.5 units/µl PKA (N=3 pigs, n=1463 events), or 0.4 µM MLCK (N=3 pigs, n=2292 events sarcomeres). Data is shown as mean ± SEM, * represents p<0.05 determined using the Student’s T-test.

**Table 1.**
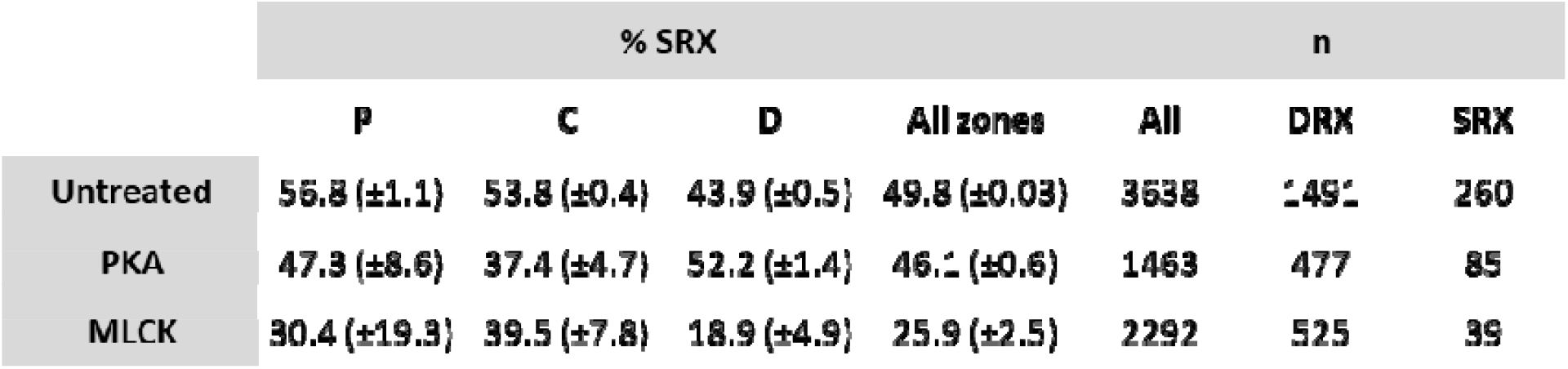
Quantification of the percentage of myosins in the SRX state in untreated porcine cardiac myofibrils (N = 3) and myofibrils treated with PKA (N = 3) or MLCK (N = 2). Percentage SRX (% SRX) was calculated as an average across the entire sarcomere (‘all zones’) and calculated within the P-, C- and D-zones of the sarcomere. Number of events (n) are provided for total tracked events and number of events attributed to DRX and SRX biochemical states. Data are mean ± SEM using n.

**Table 2.** Summary of percent change in myosin heads in the SRX state following treatment of myofibrils with PKA or MLCK relative to data from untreated myofibrils. Data are mean ± SEM (n given in Table 1).

|  | Δ SRX (%) |  |  |  |
| --- | --- | --- | --- | --- |
|  | P | C | D | All zones |
| PKA | -9.5 (±8.7) | -16.4 (±4.7) | 8.3 (±1.5) | -3.8 (±0.6) |
| MLCK | -26.4 (±19.3) | -14.3 (±7.8) | -25 (±4.9) | -23.9 (±2.5) |

Treatment with MLCK phosphorylates myosin’s RLC and is expected to reduce the population of SRX (Morano et al. 1985; Sweeney and Stull 1986; Olsson et al. 2004; Stelzer, Patel, and Moss 2006; Kampourakis, Sun, and Irving 2016). However, in a gel-based assay we were unable to detect a statistically significant increase in the phosphorylation of RLC (Fig 2D). Nevertheless, a clear reduction in the SRX population by 23.9% was observed across all zones (to 25.9% ± 2.5%, Figure 3 and Table 1). Examination of the distribution of percent SRX indicated a 26.4% reduction in the P-zone (to 30.4% ± 19.3%, Figure 3 and Table 1) and a similar reduction of 25.0% in the D-zone (to 18.9% ± 4.9%, Figure 3 and Table 1) compared to the C-zone, which showed a much smaller reduction of 14.3% (to 39.5% ± 7.8%, Figure 3 and Table 1). Combining phosphorylation from MLCK and PKA for both RLC and cMyBP-C respectively was not performed because of the extended period of incubation required for the specific conditions needed for each treatment, which leads to deterioration of the myofibrils.

## Discussion

Cardiac muscle contraction is controlled on a beat-to-beat basis by the release and sequestration of calcium ions. However, the heart also requires a more nuanced response to external factors such as load and ß-adrenergic stimulation. These affect both the timing of contraction and the force produced. Myosin motors generate force in the extremely dense environment of proteins arranged along the thick and thin filaments within cardiac myofibrils. By altering myosin’s ability to interact with actin, the level of force can be changed; this leads to the current view of thick filament regulation, which suggests that myosins can exist in a state either more (DRX) or less (SRX) likely to interact with actin. These states are characterized by their ATPase activities, with the DRX being more catalytically active than SRX. The precise conformational nature and location of these myosin states along the thick filament is not known. Using single molecule imaging, our results show that PKA phosphorylation releases heads from the lower activity state preferentially in the cMyBP-C containing C-zone of the sarcomere. Treatment with MLCK to phosphorylate myosin’s regulatory light chain led to greater myosin activation across all sarcomeric zones, although the effect was blunted in the C-zone. Together, these results provide insight into the complex interactions of myosin with its regulatory proteins in the milieu of the sarcomere. These results also suggest that RLC phosphorylation drives the activation of heads in the sentinel D-zone (Roger Craig and Padrón 2022) but, that cMyBP-C can still repress myosins in the C-zone. This presents a hierarchy of regulation, where the phosphorylation state of cMyBP-C governs the activity of myosin heads.

### cMyBP-C phosphorylation reduces myosin activity in the D-zone and activates in the C-zone

ß-adrenergic stimulation leads to the activation of PKA which has a number of known targets in the sarcomere. One of these, cMyBP-C, has three to four serine phosphorylation sites in the N-terminal M-domain (Jia et al. 2010; Ponnam et al. 2019; McNamara, Singh, and Sadayappan 2019; Doh et al. 2022; Mamidi et al. 2017; Kooij et al. 2013), which are thought to affect the interactions of cMyBP-C with both myosin and actin (Mamidi et al. 2017; McNamara, Singh, and Sadayappan 2019; Shaffer, Kensler, and Harris 2009). In this study, we report an overall net increase in myosin shifting toward the DRX state after PKA treatment. When analyzed by zone, the C-zone showed the biggest increase in DRX myosins, consistent with phosphorylation of cMyBP-C de-repressing the ATP turnover rates of myosin. This effect of PKA phosphorylation was balanced by an increase in the amount of repressed myosin in the D-zone. The origin of this increase in repressed myosins is unexpected and unknown, and is potentially suggestive that other factors such as titin phosphorylation (Krüger and Linke 2006; Hamdani, Herwig, and Linke 2017) might play a role in changing the ratio of DRX to SRX myosins within regions of the thick filament. The effect of PKA phosphorylation of titin is to reduce passive muscle tension (Yamasaki et al. 2002; Krüger and Linke 2006) through the unfolding of the elastic N2B-cardiac unique sequence (N2Bus) region of titin, proximal to the Z-disc. How this might affect the D-zone activity of myosin still remains unknown. Since we did not physically extend myofibrils during the single molecule experiment, we observed sarcomere lengths of ∼1.76 µm for both untreated and PKA-treated myofibrils; resulting in near complete overlap of the A-band. Therefore, even though the N2Bus region of titin is located towards the Z-disc, this is consistent with where the D-zone would reside in our preparations, raising the possibility of an interaction between these proteins. The resulting depression of myosin activity from PKA phosphorylation would also be consistent with promoting a reduction in passive tension.

### Myosin activation through RLC phosphorylation is cMyBP-C dependent

The myosin-associated regulatory light chain (RLC), stabilizes the lever arm of myosin to maximize force transmission (Sherwood et al. 2004; Sheikh et al. 2012), alters crossbridge detachment (Scruggs et al. 2009; Toepfer, West, and Ferenczi 2016; Diffee et al. 1996; Pulcastro et al. 2016) and affects the position of the myosin heads relative to the thick filament backbone (Colson et al. 2010; Levine et al. 1996). These effects are modulated by the state of RLC phosphorylation, with greater phosphorylation leading to more thick filament disorder (Levine et al. 1996) or movement towards the thin filament and slower crossbridge detachment (Turner et al. 2023). Phosphorylation of RLC is mediated by the cardiac specific MLCK (Chan et al. 2008; Ding et al. 2010) which maintains *in vivo* levels of approximately one phosphate per pair of RLCs on a myosin dimer (Colson et al. 2010; Scruggs et al. 2010; Chang et al. 2015). Our gel-based investigation indicated no statistically significant increase in phosphorylation after MLCK phosphorylation relative to untreated control myofibrils. However, a functional difference was identified between these two treatment groups, suggesting that the gel staining was not sensitive to the increased phosphorylation. Alternatively, it is possible that MLCK was generating pockets of more myosin dimers where RLCs on both myosin heads were phosphorylated due to the local availability of the substrate, which may cause a more pronounced effect. This speculative suggestion would be a useful future investigation, and although it has been suggested that phosphorylation of RLCs is randomly distributed (Chang et al. 2015), positive cooperativity may be more difficult to detect. However, the observed functional difference offers an exciting glimpse into how phosphorylation is managed in the sarcomere. Our results suggest that phosphorylation of RLC by MLCK resulted in a large reduction of SRX myosins in the P- and D-zones, whereas the C-zone exhibited no change in the ratio of SRX or DRX myosins upon treatment with PKA or MLCK. A recent insightful study found that when cMyBP-C was knocked out in mice, RLC phosphorylation led to more pronounced effects compared to when cMyBP-C was present (Turner et al. 2023). The data presented here supports this conclusion by showing that the effects of RLC phosphorylation are blunted in the C-zone. Therefore, it is possible that the interaction of cMyBP-C with RLC (Heling, Geeves, and Kad 2020; Ratti et al. 2011) supersedes the activating effect of RLC phosphorylation or that cMyBP-C can limit the extent of RLC phosphorylation.

## Conclusions

Post-translational modifications of sarcomeric proteins through phosphorylation and dephosphorylation have been shown to play important roles in controlling muscle contraction. In this study, we have shown that PKA exercises its effect primarily in the C-zone of porcine ventricular, β-myosin heavy chain containing, myofibrils. This is consistent with cMyBP-C acting as a target for this enzyme, however, complete de-repression of myosin activity was not seen. Using MLCK to phosphorylate RLC, we found a large, significant increase in myosins shifting to the DRX state. Interestingly, the C-zone showed no further activation of myosin that was not mediated by PKA alone. These data suggest a potential hierarchy, with cMyBP-C dominating the regulation due to RLC phosphorylation.

## Data availability

Coding of custom scripts is available here: https://github.com/Kad-Lab/Myofibril-phosphorylation

## Notes

https://github.com/Kad-Lab/Myofibril-phosphorylation

### Competing Interest Statement

The authors have declared no competing interest.

