## Supplementary Material for "Spatially resolving how phosphorylation affects ß-cardiac myosin activity in porcine myofibril sarcomeres with single molecule resolution"

Running title: Single molecule imaging of phosphorylation in β-cardiac myofibrils

**Supplementary Material**

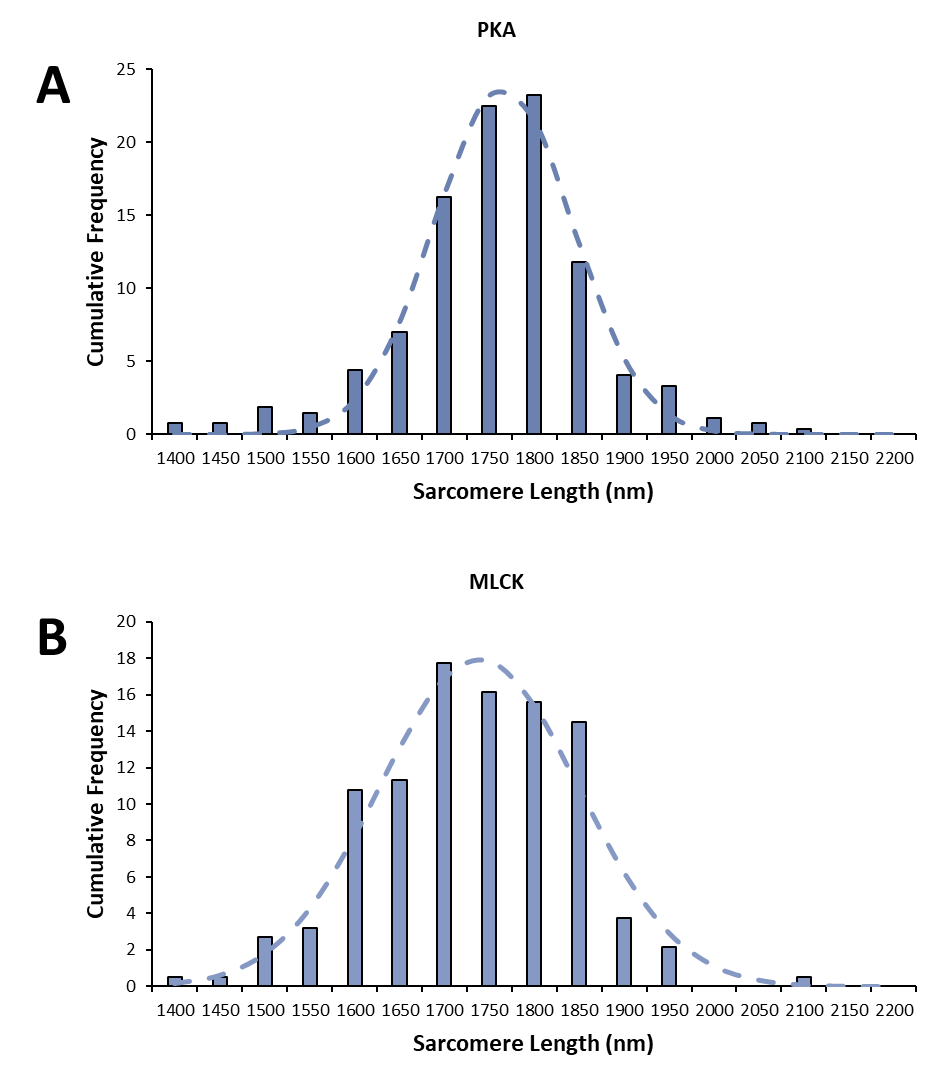

**Figure S1 Distribution of sarcomeric lengths for differentially treated cardiac myofibrils.** **A)** Histogram of the distribution of sarcomeric length for PKA treated myofibrils fitted to a Gaussian distribution (dotted line) providing a mean and SD 1766 ± 38.5 nm. **B)** MLCK treated sarcomere lengths were plotted as a histogram and fitted to a Gaussian distribution (dotted line) to give the mean and SD 1738 ± 55.6 nm.

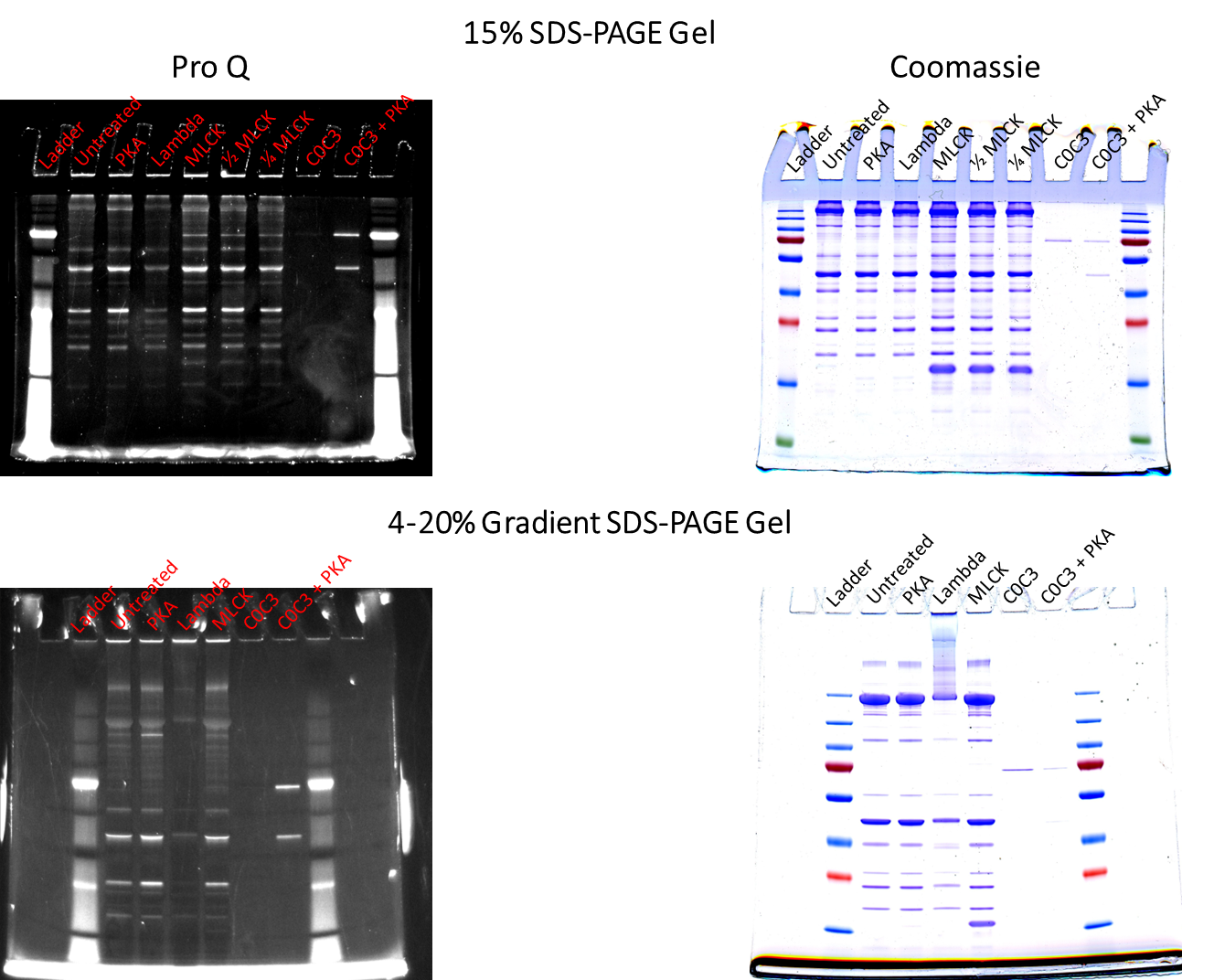

**Figure S2 Uncropped gels from Figure 2.** Both the 15% and gradient gels are shown in their entirety on the left Pro-Q-stained for phosphorylation and on the right following Coomassie staining.
